# Scalable isolation of soil genomic DNA from microbes to multicellular micro- and mesofauna

**DOI:** 10.64898/2026.01.16.699919

**Authors:** Sannakajsa Velmala, Tero Tuomivirta, Juha-Matti Pitkänen, Satu Latvala, Taina Pennanen

## Abstract

Global initiatives emphasize the need for harmonized soil biodiversity assessments. Efficient DNA extraction methods that accommodate larger soil volumes are essential for capturing higher trophic levels than bacteria and fungi and supporting extensive sampling campaigns.

We developed and evaluated a scalable, cost-efficient, and automation-ready soil DNA isolation technique alongside commercial protocols. Three starting soil amounts (0.25 g, 2.5 g, and 5 g) were tested using widely used Qiagen kits, the developed isolation method, or combinations thereof. Lysis volumes ranged from 800 µl to 15 ml, and purification employed either silica membrane or carboxyl-coated magnetic beads. Four different types of soil, both agricultural and forest soil, samples were sequenced on an Illumina MiSeq platform using universal eukaryotic primers targeting the 18S rRNA SSU region, enabling detection of non-fungal eukaryotes such as soil mesofauna and protozoa.

The developed protocol, which combined a tenfold increase in sample volume with hybrid purification steps, yielded the highest DNA recovery and consistently improved detected richness in several soil types. Species richness patterns varied by soil type and organism group: for eukaryotes and protozoa, commercial maxiprep methods along with the combination methods outperformed the miniprep approach in agricultural soils, while the developed technique excelled in coarse xeric forest soils. For metazoans, larger extraction volumes were associated with higher richness in forest soils. Our findings indicate that at least a tenfold increase in soil input compared to conventional 0.2–0.3 g is required to reliably capture mesofaunal diversity, with preliminary evidence suggesting further benefits at 20-fold volumes.

We confirm that extraction volume is a key factor shaping detection of both soil metazoan and protozoan community compositions, with effects varying by soil type and organism group. The developed scalable approach offers a practical solution for large-scale soil biodiversity assessments, aligning with global monitoring goals and enabling integration of higher trophic levels into eDNA-based frameworks.

## Introduction

Soil biodiversity underpins essential ecosystem functions, yet its comprehensive assessment remains challenging. In soil science literature, studies on “soil microbiota” have traditionally focused on bacterial communities. Even when a broader perspective is considered, research typically targets either soil microbes such as bacteria and fungi or soil fauna including animal-like protists, rarely both. This separation reflects the need for taxonomically specialized expertise and methodological differences between microbial and faunal studies. Furthermore, detecting organisms across trophic levels often requires separate sampling procedures, as larger-bodied organisms demand substantially greater soil volumes (Taberlet et al., 2012; Dopheide et al., 2019). Even single-celled eukaryotes, such as protists, are also largely absent from standard soil microbiological surveys.

Recent advances in environmental DNA (eDNA) technologies, reference databases, and bioinformatics now enable more comprehensive assessments of soil biological communities. Bottlenecks to overcome are, in turn, the great variation in the studied environment and the insufficient volumes of soil samples subjected to DNA sequencing. Reliable biodiversity monitoring requires not only adequate laboratory sample sizes but also spatially representative field sampling (Pennanen et al., 1999; Rossi & Nuutinen, 2004; Orgiazzi et al., 2018). Soil eDNA analysis has become a key tool for monitoring community composition and, in some cases, soil health. Yet, most DNA extraction protocols remain optimized for single-cell organisms.

Soil fauna play a critical role in nutrient cycling, organic matter decomposition, and soil structure. They include unicellular protistan microfauna (<0.1 mm) and multicellular mesofauna (0.1–2 mm), comprising phyla such as Arthropoda, Tardigrada, Rotifera, Nematoda, and Loricifera. These heterotrophs link microbial decomposers to higher-level consumers and enhance soil porosity through movement. As the European Soil Monitoring Law (Directorate-General for Environment, 2023) moves toward harmonised soil monitoring, there is an urgent need for standardized, cost-efficient, and scalable high-throughput DNA extraction methods. These approaches will support large-scale implementation of eDNA-based monitoring and provide robust data to maintain and restore healthy soils, in line with EU targets for 2050. Recent work by Sánchez-Cueto et al. (2025) underscores the absence of a standardized framework for soil fauna assessment using molecular approaches and pinpoints key methodological gaps that include optimized DNA extraction protocols and primers with comprehensive reference databases. Addressing these limitations is critical for advancing holistic soil biodiversity assessments that integrate trophic complexity.

Current protocols for metazoan and protozoan community analysis rely heavily on commercial kits, often using small sample sizes (0.25–0.5 g) that limit detection of larger-bodied organisms. Since 2023, studies focusing on metazoan and protozoan community analysis mainly use commercial kits. Sediment and benthic studies often rely on the DNeasy PowerMax Soil Kit (Qiagen) with 3–10 g of material (Angulo-Preckler et al., 2023; Hale et al., 2024; Han et al., 2022; Jensen et al., 2024; Maciute et al., 2025; Mazurkiewicz et al., 2024; Pawlowski et al., 2024). Some recent studies use smaller sample sizes (0.25–0.5 g) with kits such as FastDNA® SPIN kit for soil (MP Biomedicals, Santa Ana, CA, USA) (Lin et al., 2025) E.Z.N.A.® Soil DNA Kit (Omega Bio-tek, Norcross, GA, USA) (Ma et al., 2024), or PowerSoil Pro (Qiagen) kit (Xu et al., 2023). Soil metazoan studies typically use DNeasy PowerSoil Pro Qiagen kits with 0.25 g samples (Dendooven et al., 2025; He et al., 2024; Jeanbille et al., 2024; Sánchez-Cueto et al., 2025).To address the limitations of small sample sizes in commercial kits, some studies use multiple technical replicates (e.g., Dendooven et al., 2025) or pre-treatment steps such as lyophilization and homogenization. For example, Sapkota et al. (2025) lyophilized 40 g of soil and applied three consecutive bead homogenization steps to ensure complete pulverization and mixing before subsampling 0.25 g for DNA extraction. Previous comparisons indicate that 1.5–15 g of soil may be required for accurate metazoan diversity assessment (Dopheide et al., 2019), whereas high-throughput surveys such as the Land Use-Land Cover Area Frame Survey LUCAS use only 0.2 g (Labouyrie et al., 2023). This limitation emphasizes the necessity for improved, scalable protocols that accommodate larger sample volumes.

Efficient DNA extraction methods that retain microbial diversity are fundamental to advancing soil biodiversity research and contributing to European as well as global soil health initiatives. To address this gap, we aimed to develop a scalable, cost-efficient, and automation-ready high-throughput soil DNA extraction protocol capable of processing larger sample volumes and supporting extensive sampling efforts. Considering the increasing interests towards soil biodiversity monitoring, eDNA analytics that take higher trophic levels into account would be highly interesting.

## Material and Methods

### Soil material

We used soil from two agricultural fields and two types of boreal forests. The first was high mull (high organic matter >30%) agricultural soil, pH 5.7 (hereafter Amull), and the second was fine sandy till, pH 6.5 (Asandytill). The third was carex peat forest soil (Fpeat) from mesic upland forest site (vitis-idaea type and myrtillus type), pH 4.2. The fourth was coarse xeric Scots pine forest soil (CT – Calluna type) pH 4.5 (Fcoarse).

### DNA extraction methods

In short, we tested three different starting amounts of soil 0.25 g, 2.5 g, and 5 g, processed with PowerMax Soil (Qq1) or reference isolation technique DNeasy PowerSoil Pro (Qq3) (Qiagen) kits as recommended by manufacturer or with the developed novel method hereafter called the *isolation technique*, or a combination of these (Table 1). Lysis buffer volumes varied from 800 µl to 15 ml in 2 to 50 ml test tubes (Table 1). DNA binding into solid phase carrier during purification was done with binding DNA either to carboxyl-coated magnetic beads or silica membrane in the presence of NaCl and PEG or chaotropic salt, respectively. The carboxyl-coated particles used were in -house produced paramagnetic carboxyl-coated particles as described in Oberacker et al. (2019) or those commercially available from Cytiva (USA) sold under tradename Sera-Mag SpeedBeads™ Carboxylate (hydrophylic) (Table 1).The Carboxylate Sera-Mag SpeedBeads™ (hydrophylic) were washed twice with TE-buffer (10 mM Tris-HCl, 1 mM EDTA, pH 8.5) to remove sodium azide, which can interfere with downstream steps. The beads to be used were resuspended in TE buffer to a final concentration of 50 mg ml^-1^. The bound DNA was washed with 80 % ethanol and eluted in buffer C6 (Qiagen), consisting of 10 mM Tris-HCl (pH 8.5). All extractions were made in triplicates except for the Qq2 scheme which was done in duplicate (Table 1).

**Table 1.**
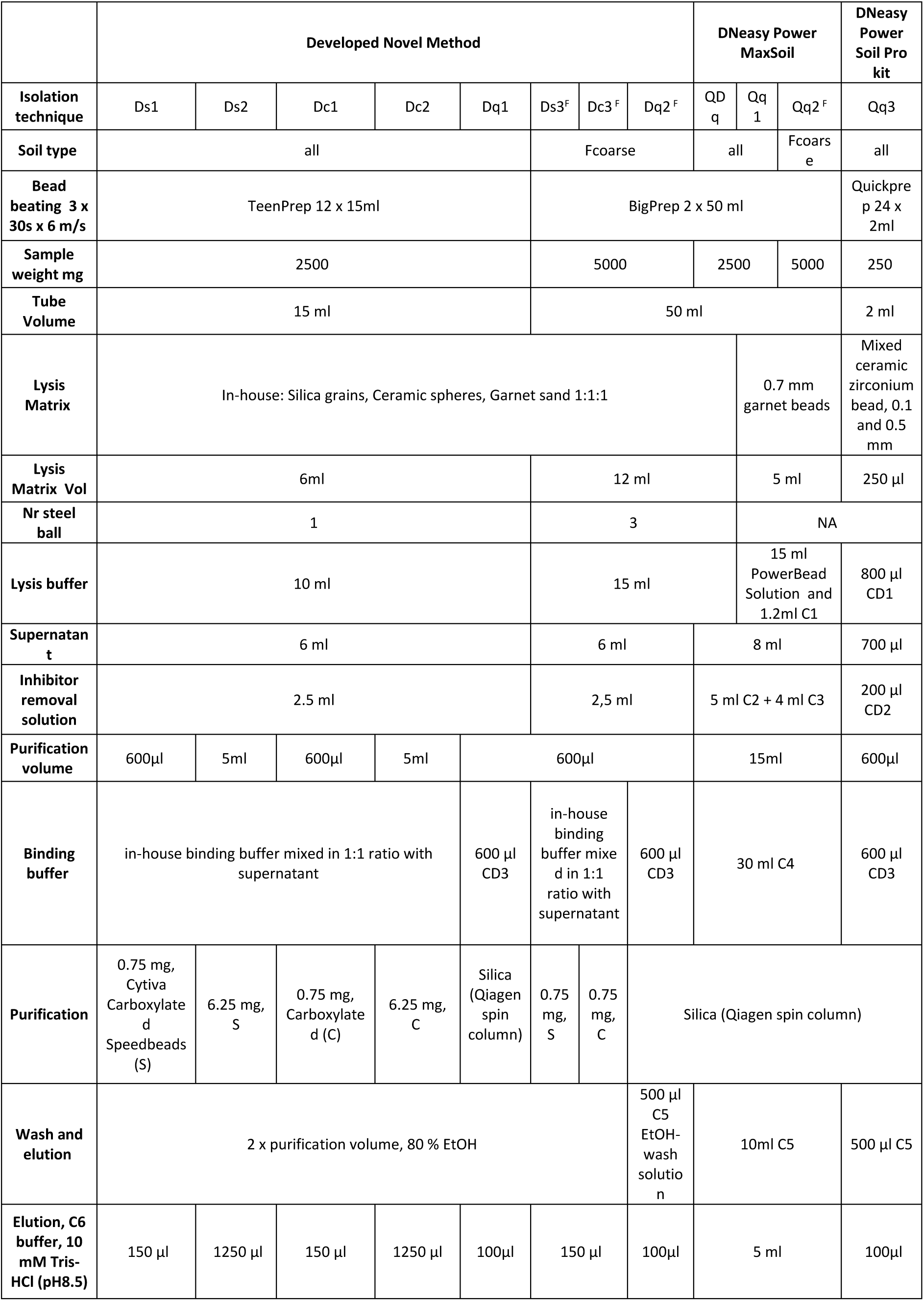
DNA isolation technique: Altogether 12 different eDNA extraction schemes and their abbreviations. ^F^ for test with large volumes we used the easiest soil type, Fcoarse.

### DNA extraction ingredients and steps

The in-house sample lysis matrix of the novel isolation technique contained equal parts (1:1:1 vol) of blasting glass spheres (Ø 0.09-0.15 mm), Zirblast® B20 ceramic spheres (Ø 0.6–0.85 mm, Saint-Gobain, France), and garnet sand (Ø 0.50–1.00 mm, JH Mining, China). In addition, one or three Gamo BB steel balls (Ø 4.5 mm, Gamo Outdoors S.L.U., Spain) were added to the lysing matrix (Table 1 and Supplementary table S3) and assembled into Ultra-high Performance 15- or 50-ml polypropylene conical test tubes (VWR International, U.S.A.). All matrix components were washed with distilled water and rinsed once with 70% EtOH before autoclaving at 121 °C for 20 minutes and oven dried at +70 °C o/n prior to tube assembly.

Sample lysis in the novel isolation technique is based on mechanical disruption of cells in the presence of a lysis buffer (Sodium Thiocyanate NaSCN 1.25 M 98% extra pure, Thermo Scientific Chemicals, U.S.A.; Disodium Phosphate (Na₂HPO₄) 0.2 M, ≥99%, GPR RECTAPUR®, VWR Chemicals, U.S.A.), with a pH of 8.4. Volumes for different isolation techniques are reported in Table 1.

Sample homogenisation in 15- and 50-ml conical test tubes was done in the FastPrep 24 machine, equipped with TeenPrep and BigPrep adapters (MP Biomedicals, U.S.A.) (Table 1). These adapters were used to ensure effective homogenisation: 3 times for 30 seconds at speed 6 m s^-1^. After homogenisation, tubes were centrifuged for 2 minutes at 16 000 x g. The samples for DNeasy PowerSoil Pro (Qiagen) method were lysed according to manufacturer’s instructions for 20 minutes in 2-ml test tubes using Vortex adapters (Vortex-Genie 2, Scientific Industries, USA).

Inhibitor removal in the developed novel isolation techniques was based on the addition of an in-house inhibitor removal solution containing aluminium chloride (AlCl_3_, 0.095 M) and ammonium acetate (NH_4_CH_3_CO_2_, 3.75 M, ≥97%, GPR RECTAPUR®, VWR Chemicals, U.S.A., pH of 6.3) to the centrifuged supernatant from the homogenisation step. Inhibitor removal was based on chemical coagulation of humic acids, proteins, and debris. To summarise, a 6-ml aliquot of the supernatant from homogenized and centrifuged lysate was pipetted into new 15-ml conical test tubes and mixed using a Vortex-Genie 2 for 5 seconds with 2.5-ml of in-house inhibitor removal solution (Table 1). Tubes were centrifuged for 3 minutes at 17 000 x g, and supernatant was moved into a new clean tube.

Following this, the DNA from the supernatant was bound to a carrier, either a carboxyl-coated particles or a silica membrane, thus facilitating its separation. In the developed novel method, DNA was bound to the carboxyl-coated particles (1.25 mg ml^-1^ of supernatant) with equal amounts of binding buffer (PEG8000 12.5%, NaCl 1.2 M, Tween20 0.05%) to the supernatant from the inhibitor removal step. The binding solution was shaken at room temperature at 1300 rpm for 5 minutes. A neodymium magnet was applied to settle all the beads to the side of the tube and the cleared solution was removed. Beads were then washed twice with equal amounts of 80% EtOH to the supernatant and shaken as described above. After each washing step, a neodymium magnet was applied to settle all the beads and ethanol was meticulously removed. Particles were incubated at room temperature for 5 minutes to dry before elution (Table 1). Elution buffer C6 (10 mM Tris-HCl and EDTA 1 mM buffer, pH 8.5) was added on to the particles, and the mixture was vortexed, and then incubated at room temperature for 5 minutes. The magnet was applied to the mixture, and the cleared supernatant (containing DNA) was collected to a fresh tube.

DNeasy PowerMax Soil (Qq1) and DNeasy Power Soil Pro (Qq3, here the reference isolation technique) kits were used as described by the manufacturer, using dedicated vortex adapters in bead beating. In addition, two additional DNeasy PowerMax Soil DNA isolation schemes were tested; One with 5 g of soil (Qq2) and the other with 2.5 g of soil (QDq) where the kit’s own lysis matrix was replaced with in-house lysis matrix with 3 steel balls (Table 1).

The quantity and quality of isolated DNA were measured using a Qubit 4 fluorometer (Thermo Fisher Scientific, U.S.A.) and a NanoDrop One Spectrophotometer (Thermo Fisher Scientific, U.S.A.), respectively. DNA was stored at -20 °C until used in downstream processes. Instructions for using the developed novel method with carboxyl-coated particles on a KingFisher™ Apex semi-automatic purification system (Thermo Fisher Scientific) can be found in the supplementary material (Tables S3 and S4). This method is adapted for 3 g soil samples in 24 deep well plates.

### Inhibition tests

To assess the quality of DNA we run qPCR inhibition tests for a subset of samples with HOT FIREPol® EvaGreen qPCR Supermix (Solis Biodyne, Estonia) which is not marketed as an inhibition resistant qPCR kit according to the manufacturer. The quality of extracted DNA as a PCR template was tested for a subsample of soils with 0.167, 0.67, and 2.67 ng/µl (low, medium, and high DNA concentration) of extracted DNA mixed with ∼10^5^ copies of target DNA (self-ligated empty cloning vector) in 6 µl reaction volume. Amplification target DNA was compared to a control reaction without soil DNA containing only target DNA (Supplementary table S2.)

### Sequencing and bioinformatics

All samples were sequenced on an Illumina MiSeq platform using universal eukaryotic primers targeting the 18S rRNA SSU region: Euk575Fngs (ASCYGYGGTAAYWCCAGC) and Euk895Rngs (TCHNHGNATTTCACCNCT), producing an amplicon of approximately 320 bp. These primers are widely used for detecting non-fungal eukaryotes, particularly soil-dwelling animals (European Commission, 2018; Guerra et al., 2021; Vasar et al., 2022). Samples were sequenced at the DNA Sequencing and Genomics Laboratory (BIDGEN), Helsinki.

Bioinformatic and statistical analyses of eDNA-based communities were performed in R (R Core Team, 2024). Raw 18S rRNA reads were trimmed with Cutadapt (Martin, 2011) and processed using a DADA2-based pipeline (Callahan et al., 2016) on the IT Center for Science (CSC), Finland’s national high-performance computing center, Puhti supercomputer. Steps included quality filtering, error learning, merging, chimera removal, and ASV inference. Taxonomy was assigned using the PR2 database v5.0.0 (Guillou et al., 2013). It is important to note, that PR2 has limited coverage and resolution for metazoans and some protozoan groups due to incomplete representation, marine bias, and inconsistent taxonomy (Guillou et al., 2013). A cladogram that illustrates sample relationships can be found in Supplementary figure S1. Phyloseq (McMurdie and Holmes, 2013), Microbiome (Lahti and Shetty, 2019), and vegan (Oksanen et al., 2020) packages were used for data analyses, and ggplot2 (Wickham, 2016) and ggpubr (Kassambara, 2025) for visualization of the data. Eukaryotes were assessed by filtering the data by the domain. The Metazoans were further assessed by selecting only Metazoa Subdivision from the Eukaryotes. Similarly, Protozoans and Protozoa like protists were filtered from the Eukaryotes by removing clades Fungi, Metazoa, Streptophyta, Archaeplastida and Gyrista for further analysis. For deeper look into the amplicon sequence abundance of Metazoa classes please see Supplementary Figure S2. For calculating observed richness, we used data agglomerated to *species level* (like OTUS) instead of single ASVs. Aggregating ASVs to the species level merges ASVs with the same species-level rank, resulting in much smaller datasets. For example, the number of ASVs for Metazoa decreased from 1994 to 165, and for Protozoa from 13471 to 623. In assessing beta diversity for ordinations and Permanova (adonis2; vegan) and pairwise comparisons pairwiseAdonis (Martinez Arbizu, 2020) we used P/A matrix instead of relative abundances and Jaccard dissimilarity. Statistically significant differences in species richness between reference isolation technique Qq3 and all other isolation techniques was tested with aov and glht with Dunnett’s post hoc comparison, multcomp (Hothorn et al., 2008).

## Results

### Isolation technique, DNA quality, yield and purity

DNA yield varied according to the soil type and the isolation technique (Supplementary table S1). In agricultural soils, the gain was low, from 3 to 14 µg/g in mull, and 14 to 38 µg/g in sandy till soil. In forest soils, the yield was higher, varying from 21 to 53 µg/g in peat, and from 15 to 38 µg/g in coarse soil. The lowest yields were obtained with Ds2 and Dc2 isolation techniques using mull soil. The highest yield in mull soil resulted with the reference isolation technique, Qq3. The highest DNA yields from forest soils were achieved with Dq1 and Ds1, and the lowest with Qq1 and QDq isolation techniques. DNA yield (µg/g soil) and DNA purity (A260/A280, A260/A230, and A340) are reported in detail in supplementary table S1 including the mean and standard deviation of the replicates.

The inhibition based on the qPCR test ranged from non-existent to complete, where no amplification of the target DNA occurred. This indicates variable DNA quality. Inhibition was not detected using the Qq3, Qq1 and Dq1 isolation techniques (see Supplementary Table S2). The highest level of inhibition occurred in schemes Dc1, Dc2 and Ds2, and amplification was either not detected or severely inhibited in reactions spiked with a high DNA concentration. Furthermore, dilution to a medium or high concentration did not improve the situation for Dc1 and Dc2 in agricultural mull soil, while for Ds2, inhibition could only be detected at the lowest DNA concentration (see Supplementary Table S2). The inhibition of forest soil DNA was much lower. Overall, silica-based purification produced higher-quality DNA than carboxyl-coated magnetic particles. However, the PCR inhibition decreased when DNA was diluted.

Based on the results above, we concluded the Carboxylate Sera-Mag SpeedBeads™ Cytiva (USA) to be more suitable for further method development than the in -house carboxyl-coated particles. A slightly modified method based on Ds1 and Ds2 isolation techniques was developed and applied also to semi-automatic scale (the program steps are presented in the Supplementary tables S3 and S4).

### Relative abundances of ASVs

The universal Eukaryote primers Euk575Fngs and Euk895Rngs resulted in amplicons that represented mainly Fungi (∼55%), Metazoa (∼15%), Cercozoa (∼10%) and Streptophyta (∼10%) (Fig. 1a). The major phyla of Metazoans were Annelida, Arachnida and Nematoda inc. class Chromadorea (Fig. 1b), and their relative abundances varied according to soil type and isolation technique. Eukaryote, metazoan and protozoan communities varied according to the sample soil, and agricultural and forest soils had markedly different community profiles. Accordingly, also the protozoan community varied according to soil type and isolation technique (Fig. 1c).

**Figure 1.**
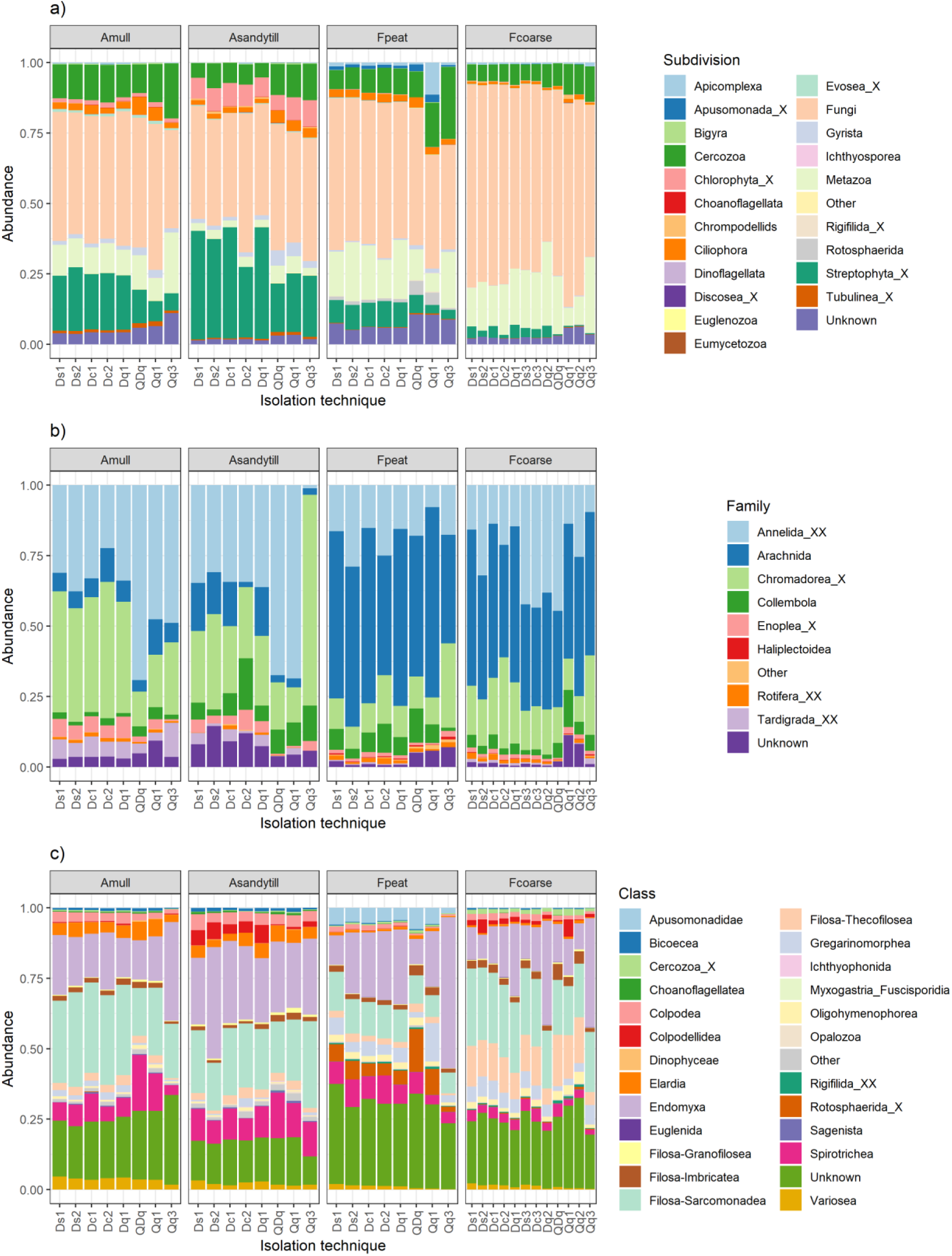
Relative abundance of a) Eukaryotes, b) Metazoans and c) Protozoans. DNA isolation technique on the x-axis. Soils include agricultural high mull (Amull) and fine sandy till (Asandytill), and forest carex peat (Fpeat) and coarse xeric soil (Fcoarse). The PR2 database uses a taxonomy that may differ from conventional classifications (Guillou et al., 2013).

### Species level richness of Eukaryotes, Metazoans and Protozoans

Species richness varied markedly among isolation techniques, with the largest deviations generally observed for the Qq3, Qq1, and Dc1, Dc2 methods compared to others (Fig. 2).

**Figure 2.**
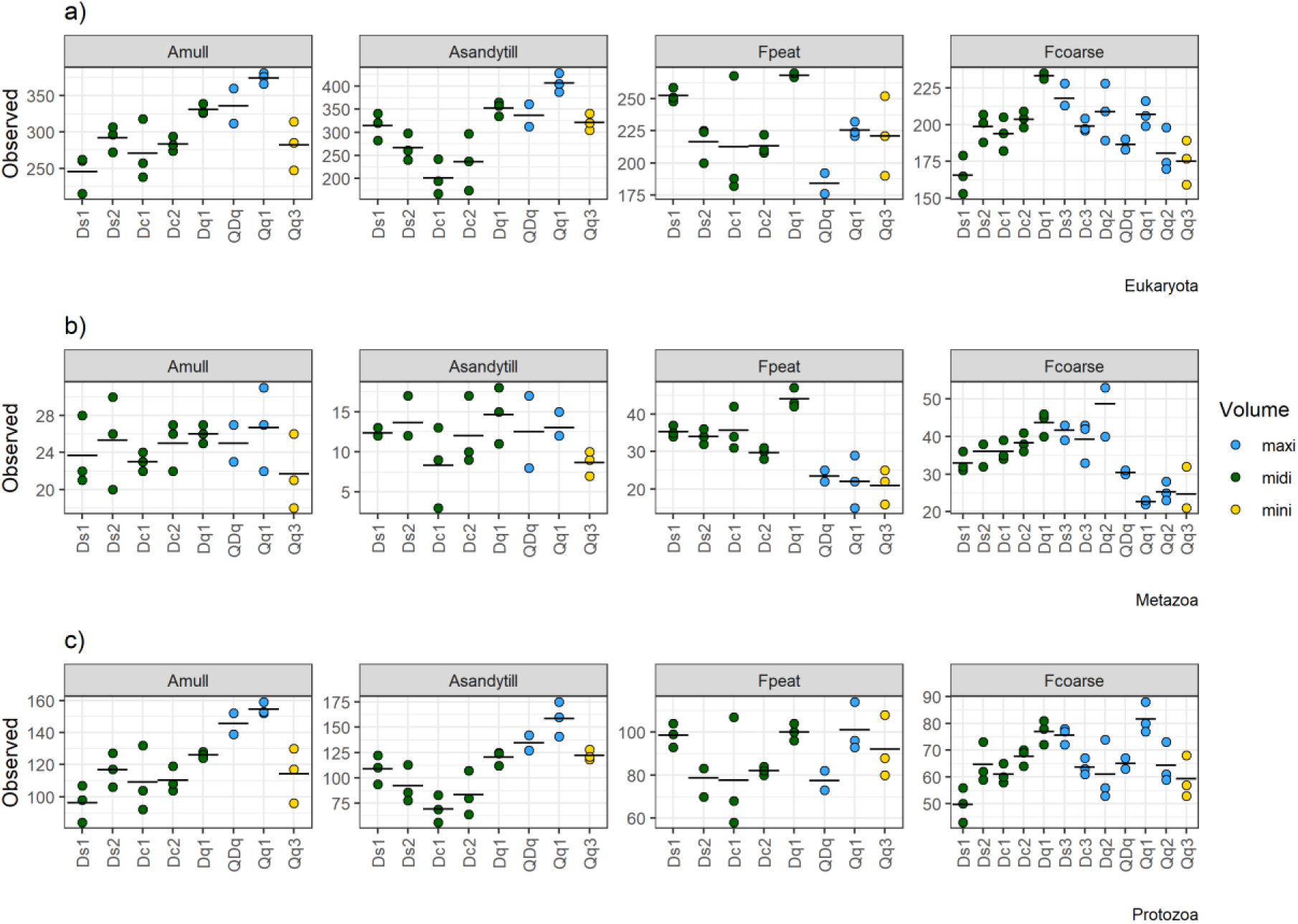
Observed species level richness of a) Eukaryotes, b) Metazoans and c) Protozoans. DNA isolation technique on the x-axis. Mean value presented as a black horizontal line. Soils include agricultural high mull (Amull) and fine sandy till (Asandytill), and forest carex peat (Fpeat) and coarse xeric soil (Fcoarse).

In agricultural soils, the eukaryote and protozoan species richness was significantly lower in the reference isolation technique Qq3 compared to the other commercial Qq1 isolation technique (p < 0.05). In sandy till soil, the Dc1 and Dc2 isolation techniques showed even lower species richness than Qq3 (p < 0.05) (Figs. 2a and 2c). However, in forest peat soil, there were no significant differences in eukaryote and protozoan species richness between Qq3 and Qq1 and all the other isolation techniques. For coarse forest soil, the developed Dq1 and Ds3 techniques exhibited significantly higher species richness than Qq3 (p < 0.001), and Dc2, Dq2, and Qq1 isolation technique also showed somewhat higher richness (p < 0.05) (Figs. 2a and 2c).

Regarding metazoan species richness, the reference isolation technique Qq3 did not differ significantly from the other techniques in agricultural soils due to high variability of replicate isolations (Fig. 2b). In contrast, in forest peat soil, Qq3 showed significantly lower metazoan richness compared to the developed Dq1 (p < 0.001), and to Dc1, Ds1, and Ds2 (p < 0.01) techniques. In coarse forest soil, metazoan richness was significantly lower in Qq3 in comparison to Dq1, Dq2, and Ds3 (p < 0.001), as well to Ds1, Dc2, Dc2, and Dc3 (p < 0.02) (Fig. 2b).

Extraction volume had a significant effect on metazoan observed richness in forest soil samples: in peat soil midi volume samples had higher richness than maxi and mini volumes (p< 0.001), and in coarse forest soil midi volume samples had higher richness than mini (p<0.05). Significant differences in species richness were observed for eukaryotes and protozoa in agricultural soil samples: mini (p<0.05, p<0.01) and midi (p<0.01, p<0.001) lower richness than maxi mull and midi lower than maxi sandy till (p<0.01) for both eukaryotes and protozoa respectively.

The annotated metazoan species showed clearly that some meso- and macrofaunal species were less found with the miniprep reference isolation technique Qq3 in comparison to the other techniques that started out with larger soil volumes. For agricultural mull soil the differences between isolation tecniques were minor, but for sandy till the Dq1 and Ds2 performed much better than the Qq3. For coarse and peat forest soil the difference was even more clear, and Dq1 revealed almost the double amount of species in comparison to the reference isolation tecnique Qq3. Moreover, all the developed in-house isolation techniques performed well, finding on average 30% more species than the Qq3. Only Qq1 had the same or smaller number of metazoan species found than Qq3 (Figure3).

**Figure 3.**
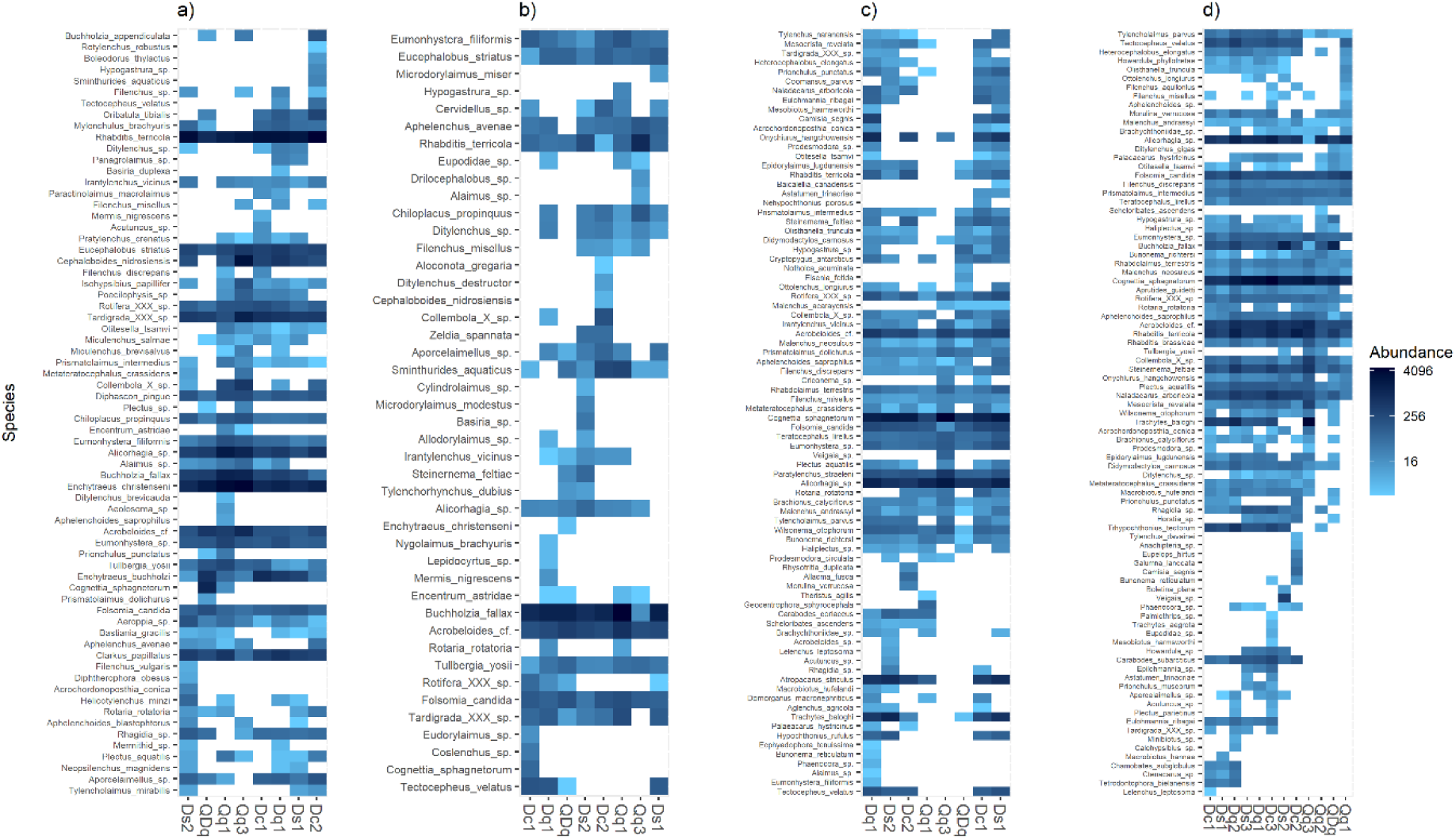
Heatmap of annotated metazoan species for each isolation technique and soiltype: a) agricultural high mull and b) fine sandy till, and c) forest carex peat and d) coarse xeric soil. Please note, that the order of the the x-axis DNA isolation technique varies according to the ordination distance matrix.

Agricultural mull soil had the lowest total richness in QDq (43 metazoan species) and highest in Dq1 (59) and Qq1 (55) (Figure 3a). Sandy till soil had the lowest total number of species with the reference isolation technique Qq3 (23) and the highest with Dq1 (36) and Ds2 (33) (Fig.3 b). Forest peat soil had the highest number of species with Dq1 and Ds1 isolation technique, and the lowest with Qq3 (Figure 3c). Coarse forest soil had the highest richness in Dq2 (83) and Dq1 (76), and the lowest in Qq1 (45) (Figure 3d). Several important enchytraeid species such as *Cognettia sphagnetorum* (forest soil:Fcoearse, Fpeat), and random occurrence also in agricultural soil) and *Enchytraeus buchholzi* (Amull), were found in samples regardless the used isolation technique. Then again in coarse forest soil some larger metazoan genera such as *Tetrodontophora sp.* were only found with isolation techniques Ds1, Dc1, Dq1, and in peat soil *Carabodes sp.* with Ds2, Dc2, Dq1 and Qq1.

### Metazoan and protozoan community composition

The NMDS ordination visualizes the dissimilarity of Metazoan and Protozoan communities in soil samples by isolation techniques (in-house isolation techniques Dq1, Ds1, Ds2 and commercial kits Qq3, Qq1, QDq). The miniprep reference isolation technique (Qq3) with low starting soil volume, 0.25 g, and the isolation techniques using in-house produced carboxyl beads (Dc) show in general larger deviation than other isolation techniques for metazoans, but the difference disappears when studying the unicellular protozoan’s and protozoa like organisms (Fig. 4, Supplementary figure 3). For both forest soil types and agricultural mull soil the coefficient of variation (CV) for observed species richness of Metazoa show that miniprep reference isolation technique Qq3 result in relatively high CV means in comparison to the developed methods, thus indicating high relative fluctuation between miniprep samples and thus inconsistent richness (Supplementary figure 3). For sandy till soil only the developed Ds1 technique showed repeatedly consistent results (Fig. 4, Supplementary figure 3).

**Figure 4.**
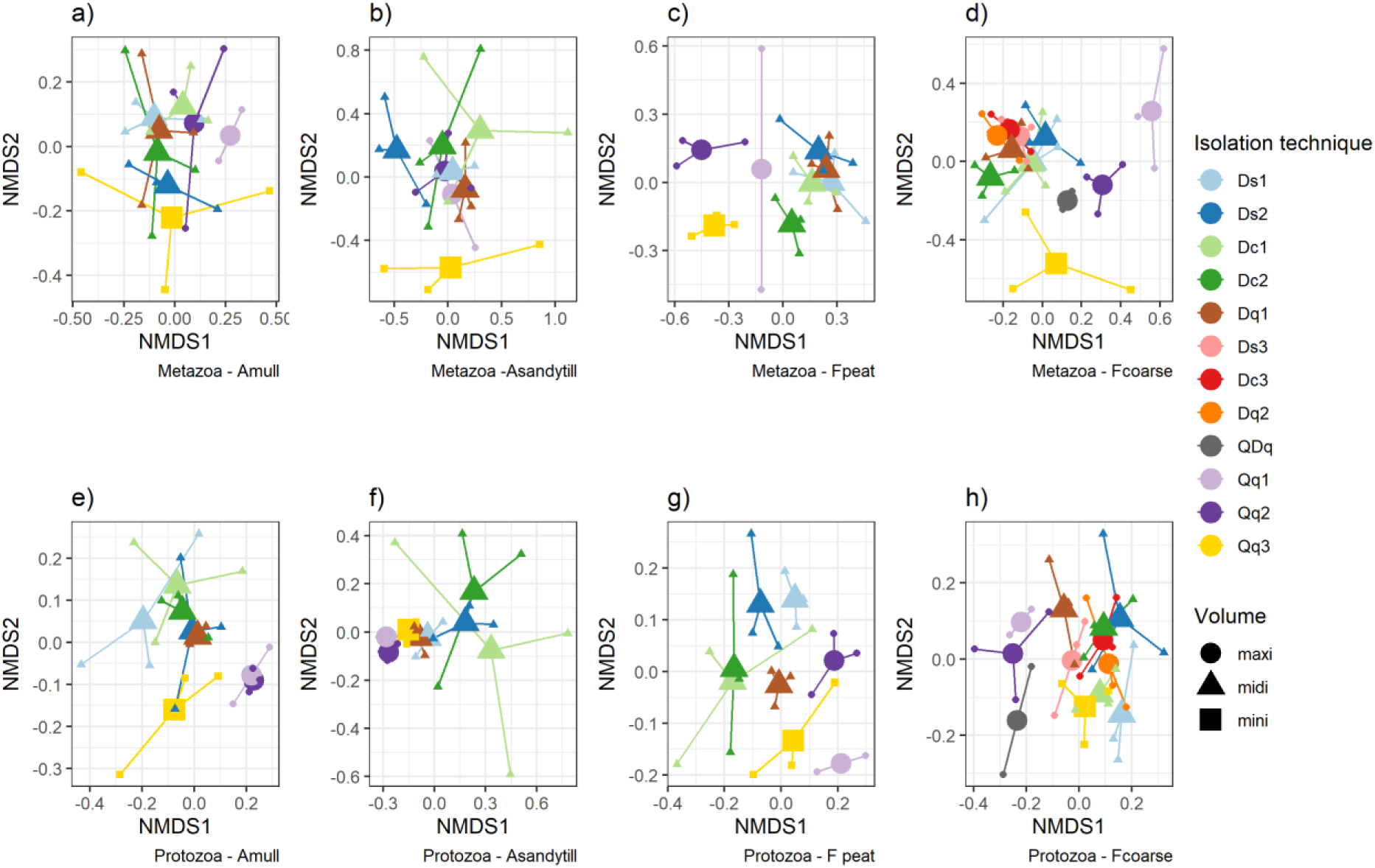
A species level 2D NMDS ordination of Metazoan communities in a) agricultural high mull (Amull) and b) fine sandy till (Asandytill), and c) forest carex peat (Fpeat) and d) coarse xeric soil (Fcoarse), and Protozoan communities e) agricultural high mull, f) fine sandy till, g) forest carex peat, h) coarse xeric soil, based on Jaccard distances calculated from presence–absence matrices. DNA isolation techniques are distinguished by colour, and extraction volumes are indicated by point shape.

The composition of soil metazoan communities was significantly influenced by extraction volume across soil types, as revealed by PERMANOVA analyses. For the agricultural mull samples (Fig. 4a), extraction volume explained a modest but significant portion of the variation in community composition (R² = 0.164, p = 0.016), with pairwise comparisons indicating small to moderate differences between mini and midi volumes (R² = 0.112, p = 0.04) and between maxi and midi volumes (R² = 0.118, p = 0.012). In contrast, no significant effects of extraction volume or isolation technique were observed for the sandy till samples (Fig. 4b). More substantial effects were detected with the forest peat samples (Fig. 4c), where extraction volume accounted for a considerable fraction of community variation (R² = 0.309, p < 0.001). Significant pairwise differences were found between mini and maxi (R² = 0.232, p = 0.037), mini and midi (R² = 0.266, p = 0.003), and maxi and midi (R² = 0.207, p < 0.001) volumes. For the coarse forest soil samples, both extraction volume (R² = 0.121, p = 0.0008) and isolation technique (R² = 0.379, p = 0.0003) contributed to explaining the community composition, with pairwise differences between mini and maxi (R² = 0.130, p = 0.021) and mini and midi (R² = 0.181, p = 0.0036) extraction volumes reflecting modest effects (Fig. 4d). However, despite the overall isolation technique effect, pairwise method comparisons were not significant after adjustment for multiple testing.

The composition of soil communities of protozoa and protozoa like organisms were also significantly influenced by extraction volume across different soil types, as demonstrated by the PERMANOVA analyses. In the mull samples (Fig. 4e), extraction volume explained a moderate portion of the variation in community composition (R² = 0.193, p < 0.001), with pairwise comparisons revealing a strong difference between mini and maxi volumes (R² = 0.294, p = 0.032) and a moderate difference between maxi and midi volumes (R² = 0.160, p < 0.001). Similarly, for the sandy till samples (Fig. 4f), both extraction volume (R² = 0.153, p = 0.011) and isolation technique (R² = 0.269, p = 0.034) explained modest but significant variation in protozoan community composition. Pairwise comparisons indicated small differences between midi and maxi volumes (R² = 0.135, p = 0.017), although, as with metazoans, pairwise isolation technique comparisons were not significant after correction for multiple testing. Extraction volume explained a moderate portion of the variation in the peat samples (Fig. 4g), where it accounted for a moderate portion of community variation (R² = 0.181, p < 0.001). Significant pairwise differences were found between mini and maxi (R² = 0.237, p = 0.040), mini and midi (R² = 0.094, p = 0.040), and maxi and midi (R² = 0.138, p < 0.001) volumes. In the coarse samples (Fig. 4h), both extraction volume (R² = 0.08, p = 0.008) and isolation technique (R² = 0.350, p < 0.001) contributed to explaining protozoan community composition, though the effect of extraction volume was limited. Notably, significant pairwise differences were detected between midi and maxi extraction volumes (R² = 0.058, p = 0.029). However, consistent with other sites, pairwise isolation technique comparisons did not remain significant after multiple testing adjustments.

## Discussion

A harmonized DNA extraction methodology is essential for accurate soil biodiversity assessments. Yet many existing methods lack scalability or compromise community representation. We present a cost-efficient, automation-ready protocol designed for high-throughput processing of larger soil volumes. This approach improves recovery of diverse microbial taxa and supports eDNA-based monitoring of micro- and mesofauna as well as unicellular eukaryotes, enabling comprehensive and reproducible biodiversity surveys that advance global soil health initiatives.

The eDNA yields based on in-house developed lysis matrix, lysis buffer, and inhibitor removal solution had different performance in different soil types compared to the reference isolation technique (Qq3, PowerSoil Pro, Qiagen), and maxiprep (Qq1, PowerMax Soil, Qiagen) DNA isolation kits. The Dq1 isolation technique, which combined the developed method with PowerSoil Pro purification steps, produced the highest DNA yield of all the isolation techniques, except for agricultural mull soil, for which the yield was slightly less than for the commercial kits. Also, the other developed isolation techniques with carboxyl-coated paramagnetic beads purification outcompeted to the reference isolation technique, Qq3, in DNA yield with peat and sandy till soil.

Carboxyl-coated paramagnetic bead isolation techniques offer easy, scalable and automatable solutions for eDNA isolation, eliminating the need for commercial kits. The main disadvantage of this method, as revealed by this study, is the lower quality of the obtained DNA, which leads to PCR inhibition. A340 values, which indicate a high concentration of particles and/or humic acids (Devi et al., 2015), were found to be between five and 150 times higher for eDNA isolated using the reference isolation technique (Qq3), depending on the isolation technique and soil type. However, PCR inhibition can be reduced by using a diluted DNA sample. Thus, all isolation schemes produced qPCR-amplifiable DNA when diluted within the limits recommended by the manufacturer (0.01 ng/µl to 4 ng/µl), as supported by our inhibition tests. To summarise, the commercial kits offered superior DNA quality over carboxyl-coated paramagnetic beads-based DNA isolation techniques described here. Similar high performance in DNA quality was observed with the Dq1 isolation technique, which combined the developed method with PowerSoil Pro purification steps where DNA was bound into a silica membrane in presence of chaotropic salt.

Humic acids are usually considered as the main PCR inhibiting substances in soil which can be removed by chemical flocculation (Braid et al., 2003). Based on the safety datasheets and patents of some commercial DNA extraction kits, AlCl_3_ is used for inhibitor removal. The similar size and charge characteristics of dsDNA and humic acids often leads to undesired co-precipitation of DNA (Wnuk et al., 2020) along with humic acids. Here we used 2.4:1 ratio between supernatant and inhibitor removal solution. However, this ratio should be optimized for every soil type especially when using carboxyl-coated paramagnetic beads. In the follow up work Saartama et al. (2026), 3:1 ratio was found to be more suitable for agricultural soil. Opposite to many commercial kits, our developed protocol uses Sodium Thiocyanate in the lysis buffer which further diminished unwanted co-extraction of humic acids into lysate / supernatant after initial homogenization. It also has lower toxicity as Guanidine Thiocyanate.

DNA isolation technique influenced species richness, but effects varied by soil type and organism group. For eukaryotes and protozoa, the reference isolation technique (Qq3, PowerSoil Pro, Qiagen) yielded lower richness than the commercial maxiprep (Qq1) in agricultural mull and sandy till soils, with some developed isolation techniques performing even worse in sandy till. In forest peat soil, isolation techniques did not differ, whereas several developed isolation techniques outperformed Qq3 in coarse forest soil, indicating higher richness with the developed approaches. For metazoans, Qq3 showed no clear differences in agricultural soils due to high variability within replicates but consistently yielded lower richness in forest peat and coarse soils compared to developed isolation techniques.

Extraction volume also mattered. Midi volumes increased metazoan richness in forest soils, while maxi volumes gave the highest eukaryote and protozoan richness in agricultural soils. Larger sample volumes are known to better capture soil heterogeneity (Dopheide et al., 2019; Sapkota et al., 2025; Taberlet et al., 2012). Consistent with Sapkopta et al. (2025), no single method captured the full diversity; isolation techniques complemented each other for metazoan detection. Our developed method lyses soil samples directly, avoiding sieving and centrifugation that may exclude macrofauna fragments or free DNA (Dopheide et al., 2019). DNA extraction sample size seems to be critical for metazoan biodiversity estimates in forest soils, though effects are non-linear. Surprisingly the developed isolation techniques outperform even commercial maxiprep kits. Unlike Dopheide et al. (2019), we also observed increased prokaryotic diversity with larger extraction volumes.

Community composition was influenced by isolation technique overall, but pairwise comparisons were not significant after correction due to low replication. However, our main focus was on the extraction volume which had a clear influence on community composition, varying by organism group and soil type. For metazoans, the mini–midi and mini–maxi contrasts were most pronounced in forest peat and coarse soils. They were minor in agricultural mull, where the differences in mini and maxi volumes were substantial, and absent in sandy till. Protozoan communities were even more sensitive, especially in forest peat and mull soils, where larger volumes improved representation despite their small size likely reflecting high diversity and spatial heterogeneity. Better detection of protozoan predators may provide insights into trophic relationships, as microbial richness contributes to soil function through complex microbiome associations (Wagg et al., 2019). Overall, these patterns highlight that scaling extraction volume can significantly alter biodiversity profiles. The combined procedure, which took advantage of lysing 10-fold sample mass but purified only a small subset of the lysate in commercial silica-based membrane, seemed to be very good in terms of DNA quality, yield and obtained richness in the tested soil types. However, for a high-throughput method, the isolation techniques using carboxyl-coated paramagnetic beads offer superior scalability at a reasonable cost.

Large variation in the body-size of soil eukaryotes from micro- to macrofauna requires attention to sampling strategy and sample size. The available commercial extraction kits are mostly designed and suitable to the small-sized microbes. In traditional morphological identification it is well established that high sample volumes are needed to cover an adequate number of individuals (Smith et al., 2008). However, in eDNA-based analytics, where the quantitativity issue is also in general very limited, the question of the portion of the truly captured diversity is easily neglected with these bigger and more sparsely occurring organisms (Kinzinger et al., 2023). Our study did not address sampling size considerations for macrofauna; rather, it focused on scaling soil sample volumes within the range appropriate for high-throughput DNA extraction targeting micro- and mesofaunal communities, alongside developing cost-effective purification protocols compatible with standard laboratory infrastructure. Nevertheless, on the contrary to the recent monitoring using smaller sample size and varying soil layers (Köninger et al., 2023), our small test set of the most common soil types indicated a higher richness of mesofauna in forest soils than in the two more intensively managed agricultural soils. We therefore propose including mesofauna in soil microbial analyses as it encompasses keystone species in the decomposer system, such as the omnivorous enchytraeid *Cognettia sphagnetorum* (Huhta et al., 1998).

Together, these findings underscore that extraction volume is a key factor shaping detection of both soil metazoan and protozoan community composition, with effects varying by soil type and organism group. In line with our expectations, the effect of analyzed soil volume was bigger for meso- and macrofaunal than protozoan communities. While isolation technique effects were detected in some cases, their influence appears more subtle and requires cautious interpretation. Nevertheless, our study highlights the importance of carefully selecting extraction protocols tailored to specific soil environments and target communities to accurately capture soil biodiversity patterns. DNA extraction method can substantially affect richness estimates, particularly in coarse and organic-rich soils, and that reliance on a single small-scale method may underestimate soil biodiversity. Improved coverage, high representativeness and cost-efficiency of eDNA-based assessments will be increasingly important tool to be considered due to the increasing needs for soil monitoring (FAO, 2024).

## Data availability

Data available upon publication from 10.5281/zenodo.15771097 and raw sequence data are deposited in the sequence read archive SRA of the NCBI database **SUB14139519**

## Acknowledgements

We thank Tuija Hytönen for technical assistance in the laboratory and Marleena Hagner for her expertise and help with identification and taxonomy of the soil invertebrate communities.

## Author contributions

Conceptualization (SV ,TT, TP, JMP, SL); Data curation (SV, TT, JMP); Formal analysis (SV); Funding acquisition (SV, TP); Investigation (SV, TT, JMP, SV); Methodology (TT, JMP, TP, SV); Project administration (SV); Resources (SV); Validation (JMP); Visualization (SV); Roles/Writing - original draft (SV, TT, TP); and Writing - review & editing (JMP, SL).

