## Supplementary Information for "Scalable isolation of soil genomic DNA from microbes to multicellular micro- and mesofauna"

**Supplementary material to the publication** *Scalable isolation of soil genomic DNA from microbes to multicellular micro‑ and mesofauna*

**Table of content**

DNA yield

Supplementary Table S1. DNA quality metrics across soil samples.	2

DNA quality

Inhibition tests	3

Supplementary table S2. Results of inhibition test.	3

High throughput DNA extraction with King Fisher apex

Supplementary table S3 Used buffers and lysis matrix.	4

Supplementary table S4 KingFisher™ Apex	system program steps.	4

Amplicon sequencing results

Supplementary table S5. Summary of amplicon sequencing.	5

Supplementary figure S1. Cladogram on clustering of soil samples.	5

Supplementary figure S2. Sequence abundance of Metazoan classes.	6

Supplementary figure S3. Coefficient of variation on Metazoan species richness	6

### DNA yield

**Supplementary table S1.** DNA quality metrics across soil samples, including i) DNA yield (µg g⁻¹ soil; mean ± SD), ii) individual replicate yields, and spectrophotometric purity ratios iii) A260/A280, iv) A260/A230 together with v) A340 measurements indicating potential background absorbance. NA = not assessed. The samples used in the qPCR inhibition tests (table S2) are highlighted in bold.

|  |  | AMull | | | | | ASandyTill | | | | | Fpeat | | | | | Fcoarse | | | | |
| --- | --- | --- | --- | --- | --- | --- | --- | --- | --- | --- | --- | --- | --- | --- | --- | --- | --- | --- | --- | --- | --- |
| Isol. tech | Repl. | i | ii | iii | iv | v | i | ii | iii | iv | v | i | ii | iii | iv | v | i | ii | iii | iv | v |
| Ds1 | a | 4.3±0.9 | 5.3 | 1.6 | 0.9 | 0.3 | 36.7±7.3 | 37.9 | 1.6 | 1.0 | 1.6 | 50.1±3.9 | **50.1** | **1.6** | **0.9** | **2.4** | 50.4±3.5 | **46.5** | **1.8** | **1.6** | **0.4** |
|  | b |  | **4.0** | **1.6** | **0.8** | **0.3** |  | 28.9 | 1.7 | 1.1 | 1.5 |  | 54.0 | 1.6 | 0.9 | 3.3 |  | 53.3 | 1.8 | 1.6 | 0.4 |
|  | c |  | 3.5 | 1.5 | 0.8 | 0.2 |  | 43.3 | 1.6 | 0.9 | 2.5 |  | 46.2 | 1.6 | 1.0 | 2.3 |  | 51.4 | 1.8 | 1.5 | 0.5 |
| Ds2 | a | 3±0.9 | **2.0** | **1.5** | **0.7** | **0.6** | 36.3±1.8 | 35.3 | 1.7 | 1.1 | 2.4 | 42.2±7.4 | **33.9** | **1.7** | **1.2** | **1.4** | 35±5.4 | **29.6** | **1.8** | **1.6** | **0.6** |
|  | b |  | 3.9 | 1.6 | 0.9 | 0.5 |  | 35.2 | 1.7 | 1.2 | 1.4 |  | 44.5 | 1.7 | 1.2 | 1.3 |  | 40.4 | 1.9 | 1.7 | 0.5 |
|  | c |  | 3.1 | 1.6 | 0.8 | 0.5 |  | 38.4 | 1.7 | 1.1 | 1.3 |  | 48.3 | 1.7 | 1.2 | 1.5 |  | 35.2 | 1.9 | 1.6 | 0.7 |
| Dc1 | a | 3.8±0.7 | 4.1 | 1.6 | 0.7 | 16.8 | 35.6±5.2 | 41.6 | 1.6 | 0.8 | 14.3 | 43.9±5.2 | **45.1** | **1.7** | **1.0** | **3.0** | 38±1.1 | **38.8** | **1.7** | **0.9** | **4.7** |
|  | b |  | **3.0** | **1.5** | **0.7** | **15.4** |  | 32.2 | 1.6 | 0.8 | 12.8 |  | 48.3 | 1.7 | 0.9 | 3.0 |  | 38.3 | 1.7 | 0.9 | 4.5 |
|  | c |  | 4.2 | 1.5 | 0.7 | 14.2 |  | 33.0 | 1.6 | 0.8 | 12.6 |  | 38.2 | 1.7 | 97.0 | 2.8 |  | 36.7 | 1.7 | 0.9 | 5.1 |
| Dc2 | a | 3.6±1.4 | **2.0** | **1.5** | **0.6** | **5.3** | 38.5±2.3 | 39.6 | 1.6 | 0.9 | 9.2 | 49.8±2.3 | **47.3** | **1.7** | **0.8** | **4.4** | 42.7±0.4 | **43.2** | **1.7** | **0.9** | **4.3** |
|  | b |  | 4.1 | 1.5 | 0.6 | 5.6 |  | 35.8 | 1.6 | 0.8 | 8.1 |  | 50.1 | 1.7 | 0.8 | 4.5 |  | 42.6 | 1.7 | 0.8 | 7.1 |
|  | c |  | 4.7 | 1.5 | 0.7 | 8.7 |  | 40.1 | 1.7 | 0.8 | 6.3 |  | 51.8 | 1.7 | 0.8 | 5.7 |  | 42.4 | 1.7 | 0.8 | 6.2 |
| Dq1 | a | 5.2±1.3 | 6.6 | 1.8 | 0.6 | 0.1 | 38.2±2.7 | 39.1 | 1.8 | 2.2 | 0.2 | 53.6±6.4 | **51.0** | **1.8** | **2.9** | **0.1** | 46.9±0.4 | **47.0** | **1.9** | **1.1** | **0.1** |
|  | b |  | **5.0** | **1.8** | **0.4** | **0.1** |  | 35.1 | 1.8 | 0.9 | 0.2 |  | 60.9 | 1.8 | 2.1 | 0.2 |  | 47.3 | 1.9 | 1.1 | 0.0 |
|  | c |  | 4.0 | 1.9 | 0.1 | 0.0 |  | 40.3 | 1.8 | 1.2 | 0.3 |  | 49.0 | 1.9 | 2.4 | 0.1 |  | 46.5 | 1.9 | 1.8 | 0.0 |
| Ds3 | a |  | NA | NA | NA | NA |  | NA | NA | NA | NA |  | NA | NA | NA | NA | 27.6±0.5 | **28.1** | **1.8** | **1.6** | **0.6** |
|  | b |  | NA | NA | NA | NA |  | NA | NA | NA | NA |  | NA | NA | NA | NA |  | 27.5 | 1.9 | 1.7 | 0.3 |
|  | c |  | NA | NA | NA | NA |  | NA | NA | NA | NA |  | NA | NA | NA | NA |  | 27.1 | 1.9 | 1.7 | 0.4 |
| Dc3 | a |  | NA | NA | NA | NA |  | NA | NA | NA | NA |  | NA | NA | NA | NA | 26.1±1.9 | **23.9** | **1.8** | **1.2** | **2.2** |
|  | b |  | NA | NA | NA | NA |  | NA | NA | NA | NA |  | NA | NA | NA | NA |  | 26.7 | 1.8 | 1.2 | 2.5 |
|  | c |  | NA | NA | NA | NA |  | NA | NA | NA | NA |  | NA | NA | NA | NA |  | 27.6 | 1.8 | 1.2 | 2.7 |
| Dq2 | a |  | NA | NA | NA | NA |  | NA | NA | NA | NA |  | NA | NA | NA | NA | 26.2±1.2 | **24.9** | **1.9** | **1.1** | **0.0** |
|  | b |  | NA | NA | NA | NA |  | NA | NA | NA | NA |  | NA | NA | NA | NA |  | 27.0 | 1.9 | 1.4 | 0.0 |
|  | c |  | NA | NA | NA | NA |  | NA | NA | NA | NA |  | NA | NA | NA | NA |  | 26.6 | 1.9 | 1.4 | 0.0 |
| QDq | a | 8.6±1.2 | 7.8 | 1.9 | 1.1 | 0.1 | 13.6±0.1 | 13.7 | 1.8 | 1.3 | 0.1 | 21±3.7 | **23.6** | **1.6** | **1.0** | **0.3** | 25±5.3 | **28.7** | **1.8** | **1.3** | **0.1** |
|  | b |  | **9.4** | **1.8** | **1.2** | **0.1** |  | 13.6 | 1.8 | 1.3 | 0.1 |  | 18.4 | 1.7 | 1.0 | 0.2 |  | 21.2 | 1.7 | 1.3 | 0.1 |
| Qq1 | a | 10±1.4 | **8.9** | **1.7** | **1.5** | **0.1** | 16.6±1.4 | 16.2 | 1.8 | 1.6 | 0.1 | 24.9±6.2 | **18.4** | **1.6** | **1.5** | **0.1** | 25±5.5 | **19.1** | **1.8** | **1.6** | **0.1** |
|  | b |  | 11.5 | 1.8 | 1.3 | 0.1 |  | 18.1 | 1.8 | 1.6 | 0.1 |  | 30.7 | 1.7 | 1.5 | 0.1 |  | 30.0 | 1.7 | 1.2 | 0.1 |
|  | c |  | 9.5 | 1.7 | 1.7 | 0.1 |  | 15.4 | 1.9 | 1.8 | 0.1 |  | 25.7 | 1.7 | 1.5 | 0.1 |  | 25.8 | 1.7 | 1.5 | 0.1 |
| Qq2 | a |  | NA | NA | NA | NA |  | NA | NA | NA | NA |  | NA | NA | NA | NA | 14.9±1.3 | **13.8** | **1.7** | **1.2** | **0.1** |
|  | b |  | NA | NA | NA | NA |  | NA | NA | NA | NA |  | NA | NA | NA | NA |  | 16.4 | 1.7 | 1.2 | 0.2 |
|  | c |  | NA | NA | NA | NA |  | NA | NA | NA | NA |  | NA | NA | NA | NA |  | 14.4 | 1.7 | 1.7 | 0.1 |
| Qq3 | a | 13.6±3.7 | **9.5** | **1.8** | **0.5** | **0.1** | 30.8±3.5 | 27.0 | 1.9 | 0.9 | 0.1 | 35±5 | 40.5 | 1.8 | 1.5 | 0.1 | 44.3±6.6 | **37.2** | **1.9** | **1.4** | **0.0** |
|  | b |  | 14.6 | 1.9 | 0.7 | 0.1 |  | 33.8 | 1.9 | 1.3 | 0.1 |  | **33.9** | **1.8** | **1.5** | **0.1** |  | 50.1 | 1.9 | 1.7 | 0.0 |
|  | c |  | 16.6 | 1.8 | 0.6 | 0.1 |  | 31.7 | 1.8 | 1.3 | 0.1 |  | 30.7 | 1.8 | 1.4 | 0.1 |  | 45.7 | 1.9 | 1.6 | 0.1 |

### DNA quality

#### Inhibition tests

The quality of DNA to be used as PCR template was tested by inhibition test containing 0.167, 0.67, or 2.67 ng /µL of extracted DNA mixed with approximately 10^5^ copies of target DNA (empty, self-ligated pGEM-T easy; Promega, U.S.A.) and compare amplification to a reaction without extracted soil DNA in a 6 µl qPCR-reaction. qPCR reactions contained also 1 x HOT FIREPOL EvaGreen qPCR Supermix (Solis Biodyne, Estonia) and 0.375 µM of SP6 and T7 primers. Reactions were set up with two technical replicates with MYRA liquid handling system (Bio Molecular Systems, Australia) and run in Mic qPCR cycler (Bio Molecular Systems, Australia). Cycling conditions for 40 cycles were as follows: Initial denaturation 95 ^o^C for 12 minutes, denaturation 95 ^o^C for 15 seconds, annealing 50 ^o^C for 30 seconds and extensions 72 ^o^C for 60 seconds. Fluorescence was measured from channel green at the end of each extension step to measure the buildup of multiple cloning site of pGEM-T easy vector. Inhibition was calculated as PCR efficiency (Sample + pGEM-T easy) / PCR efficiency (water + pGEM-T easy). Detection of baseline in qPCR was set to “dynamic” in BMS Workbench v1.4.5 software (Bio Molecular Systems, Australia).

**Supplementary table S2.** Results of inhibition test containing 2.67, 0.67, or 0.167 ng /µL of extracted soil DNA in a 6 µL qPCR reaction. Inhibition was calculated as PCR efficiency (Sample + pGEM-T easy) / PCR efficiency (water + pGEM-T easy). PCR efficiency values >0.95 indicate no or only mild inhibition. NA=not assessed. Inhib. no amplification.

|  |  |  | Amull (ng/µL) | |  | Fpeat (ng/µL) | | | Fcoarse (ng/µL) | | |
| --- | --- | --- | --- | --- | --- | --- | --- | --- | --- | --- | --- |
| **Isolation technique** | | **Purification** | **2.67** | **0.67** | **0.167** | **2.67** | **0.67** | **0.167** | **2.67** | **0.67** | **0.167** |
| Ds1 | DevMethod | cytiva carboxyl | 0.71 | 0.99 | 0.99 | 0.76 | 1 | 1.02 | 1 | 1.03 | 1 |
| Ds2 | DevMethod | cytiva carboxyl | Inhib. | 0.9 | 0.93 | 0.88 | 1.05 | 0.93 | 0.95 | 1.01 | 0.95 |
| Dc1 | DevMethod | own carboxyl | Inhib. | Inhib. | 0.7 | 0.79 | 1 | 0.99 | 0.89 | 1.04 | 1.03 |
| Dc2 | DevMethod | own carboxyl | Inhib. | Inhib. | 0.69 | Inhib. | 0.98 | 1.02 | 0.75 | 1.02 | 0.99 |
| Dq1 | DevMethod | silica | 0.97 | 1.02 | 1.02 | 0.96 | 1.04 | 1.04 | 1.01 | 1.03 | 1 |
| Ds3 | DevMethod | cytiva carboxyl | NA | NA | NA | NA | NA | NA | 1.03 | 1.05 | 1.01 |
| Dc3 | DevMethod | own carboxyl | NA | NA | NA | NA | NA | NA | 1.01 | 1.04 | NA |
| Dq2 | DevMethod | silica | NA | NA | NA | NA | NA | NA | 1 | 1.01 | 1.03 |
| QDq | MaxSoil | silica | 0.95 | 1.02 | 0.99 | Inhib. | 0.98 | 0.94 | 0.99 | 0.98 | 1.02 |
| Qq1 | MaxSoil | silica | 0.96 | 0.98 | 1.01 | 0.95 | 0.99 | 1.01 | 0.99 | 1.04 | 1.03 |
| Qq2 | MaxSoil | silica | NA | NA | NA | NA | NA | NA | 0.82 | 0.98 | 1.03 |
| Qq3 | SoilPro | silica | 0.96 | 1.04 | 1.04 | 0.97 | 1.04 | 1.02 | 0.98 | 1.02 | 1.02 |

### High throughput DNA extraction with King Fisher apex

Three grams of soil were placed in a 15 ml Polypropylene Co-Polymer (PPC) tube containing the lysis matrix (Table S3), followed by 10 ml of lysis buffer at room temperature. The mixture was homogenized using a Bead Ruptor Elite Bead Mill (OMNI) 3 x 30 s at 6.0 m/s, then centrifuged for 8 min at 7500 × g. Then 6 ml of supernatant was transferred to a clean 15 ml polypropylene conical tube and combined with 2 ml precipitation buffer (3:1 ratio) using a Vortex-Genie 2 for 10 s. After centrifugation for 8 min at 7500 × g, two 1.8 ml subsamples of the lysate were placed in 2 ml Eppendorf tubes and stored at −20 °C until DNA extraction. DNA was isolated from 1.8 ml lysate subsamples on 24 deep-well plates on KingFisher™ Apex (Thermo Fisher Scientific) (Table S4). The lysis, precipitation, binding, TE and wash buffer were prepared in autoclaved borosilicate glass bottles (Schott Duran & VWR) in autoclaved double-distilled water (DDW) as follows.

#### **Supplementary Table S3.** Used buffers and lysis matrix.

|  | **Ingredients** | **Notes** |
| --- | --- | --- |
| **Buffers** |  |  |
| Lysis buffer | Sodium Thiocyanate (NaSCN) 1.25 M | pH 8.5, stored at RT |
|  | Disodium Phosphate (Na2HPO4) 0.2 M |  |
| Precipitation buffer, pH 6.3 | Aluminium Chloride (AlCl3) 0.095 M | Prepare in a fume hood to avoid hydrochloride gas production. |
|  | Ammonium Acetate (NH4CH3CO2) 3.75 M | pH was adjusted to 6.3 with 1 M NaOH, stored at 4 °C |
| Binding buffer | PEG8000 12.5% | stored at RT |
|  | NaCl 1.2 M |  |
|  | Tween20 0.05% |  |
| TE buffer | Tris-HCl 10 mM | stored at RT |
|  | EDTA 1 mM |  |
| Wash buffer | EtOH 80% | 1.8 ml per well stored at RT |
| **Lysis matrix** 15 ml PPC tube  filled with 2 ml of each material | Silica grains 0.1 mm silica spheres (Saint-Gobain) Ø 0.1 mm Silicon oxide) | Rinsed with DDW to remove dust then rinsed once with 70% EtOH. Dried and sterilised o/n in an oven at 200 °C.  Steel balls rinsed in EtOH and autoclaved at 120 °C |
|  | Ceramic spheres 0.4 mm ceramic sphere (Saint-Gobain) Zirconium, Ø 0.6–0.85 mm (ZrO²: 62  %, SiO²: 28 %) |  |
|  | Garnet sand Garnet 0.50–1.00 mm |  |
|  | Stainless steel ball, 4.5 mm Ø(Gamo BB) |  |

#### **Supplementary Table S4.** KingFisher™ Apex semi-automatic purification system program steps

| **24 deep-well plates** | **Volume per well** | **Ingredients** | **Method** |
| --- | --- | --- | --- |
| **Bind** Plate 1 | 1.8 ml | Sample lysate | Mix lysate, beads, and binding buffer thoroughly.  Incubate 5–10 min at room temperature for DNA adsorption.  KingFisher   - Release 30 s, bottom mix - Mixing 4 min 30 s, half mix - Collect beads 3 x 25 s |
|  | 1.8 ml | Binding buffer |  |
|  | 45 µl | Sera-Mag™ Carboxylate-Modified Magnetic Beads (50 mg/ml) in TE-buffer. Washed twice with TE buffer to remove sodium azide buffer |  |
| **Wash Plates** Plates 2 & 3 | 1.8 ml | Ethanol 80% | KingFisher   - Release 10 s - Mixing 1 min, half mix - Collect beads 3 x 1 s, bottom mix   Repeat for Wash Plate 3 |
| **Dry** |  |  | 5 min to remove residual ethanol |
| **Elute** Elution plate, Plate 4 | 300 µl | TE buffer (Table Sxx) | KingFisher   - Elute 15 s, bottom mixing - Mixing 45 s, medium mix - Mixing 15 s, bottom mix - Collect beads 3 x 30 s   Store tubes frozen -20 °C or colder |

### Amplicon sequencing results

**Supplementary table S5**. Summary of amplicon sequencing results. Input and output of dada2 pipeline

| Sample | input raw reads | merged nonchimeric reads | ASV All eukaryotes | ASV Metazoa |
| --- | --- | --- | --- | --- |
| Sequence Blanck | 14743 | 32 | 1 | 0 |
| Ds1 | 1369239 | 789483 | 763 | 64 |
| Ds2 | 1197184 | 682088 | 747 | 64 |
| Dc1 | 1196167 | 719723 | 664 | 60 |
| Dc2 | 1136198 | 723196 | 688 | 60 |
| Ds3 | 432331 | 240842 | 815 | 107 |
| Dc3 | 306462 | 214782 | 718 | 98 |
| Dq2 | 437233 | 286283 | 777 | 116 |
| Dq1 | 2080476 | 1040820 | 970 | 83 |
| Qq1 | 1948669 | 1064139 | 1234 | 75 |
| Qq2 | 430071 | 193933 | 809 | 79 |
| QDq | 1204730 | 483562 | 945 | 75 |
| Qq3 | 2324672 | 894493 | 878 | 60 |

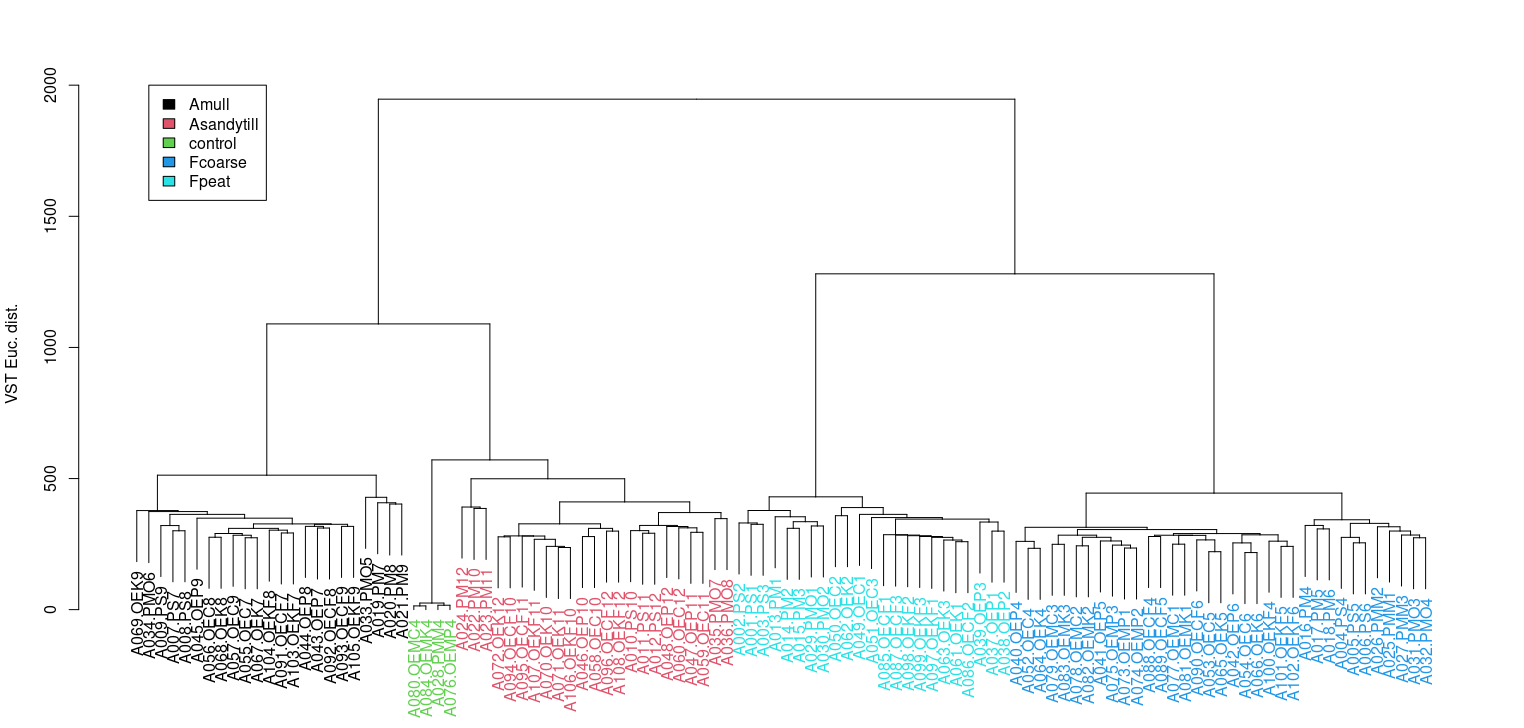

**Supplementary figure S1.** Cladogram showing clustering of soil sample types. Forest and agricultural soil samples differ greatly in their eukaryote communities. All sample types cluster together regardless of the isolation technique. Control indicate DNA extraction blank samples.

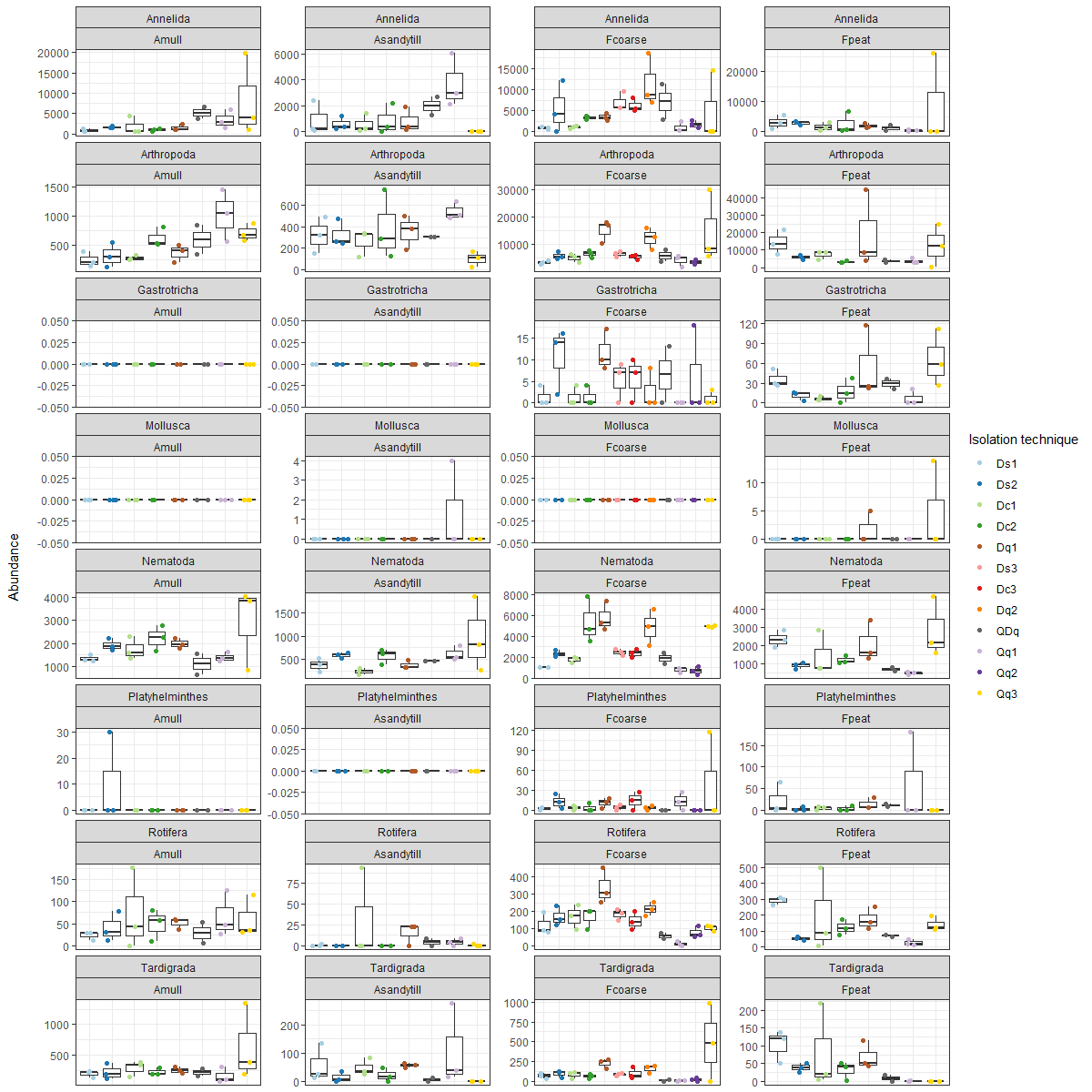

#### **Supplementary figure S2.** Sequence abundance of Metazoa classes

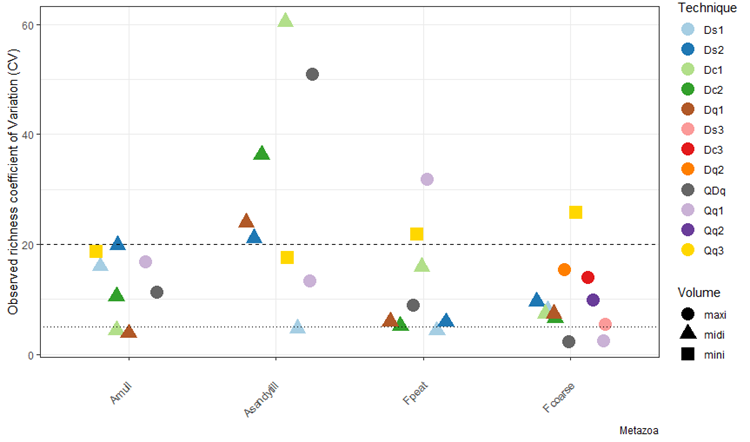

#### **Supplementary figure S3.** Coefficient of variation (CV) for observed species richness of Metazoa across different soil DNA extraction methods, extraction volumes and soil types.
